# Undersampling and the inference of coevolution in proteins

**DOI:** 10.1101/2021.04.22.441025

**Authors:** Yaakov Kleeorin, William P. Russ, Olivier Rivoire, Rama Ranganathan

## Abstract

Protein structure, function, and evolution depend on local and collective epistatic interactions between amino acids. A powerful approach to defining these interactions is to construct models of couplings between amino acids that reproduce the empirical statistics (frequencies and correlations) observed in sequences comprising a protein family. The top couplings are then interpreted. Here, we show that as currently implemented, this inference is always biased, a problem that fundamentally arises from the distinct scales at which epistasis occurs in proteins in the context of limited sampling. We show that these issues explain the ability of current approaches to predict tertiary contacts between amino acids and the inability to obviously expose larger networks of functionally-relevant, collectively evolving residues called sectors. This work provides a necessary foundation for more deeply understanding and improving evolution-based models of proteins.

## Introduction

The basic characteristics of natural proteins are the ability to fold into compact three-dimensional structures, to carry out chemical reactions, and to adapt as conditions of selection fluctuate. To understand how these properties are encoded in the amino acid sequence, a powerful approach is statistical inference from datasets of homologous sequences – the study of evolutionary constraints on and between amino acids. In different implementations, this approach has led to the successful prediction of protein tertiary structure contacts^1–4^, protein-protein interactions^5–7^, mutational effects^8–11^, and even the design of synthetic proteins that fold and function in a manner indistinguishable from their natural counterparts^12–14^. A major result from these studies is the sufficiency of pairwise correlations in multiple sequence alignments to specify many key aspects of proteins. This result motivates the search for statistical models of protein sequences that capture these correlations as a route to understanding and designing proteins.

What characteristics underlie a “good” statistical model of protein sequences? The native state of a protein represents a fine balance of large opposing forces between atoms that operate with strong distance dependence to produce marginally stable structures. Thus, many complex and non-intuitive patterns of interdependence between amino acids (epistasis) are possible, all consistent with the compact, well-packed character of tertiary structures. Indeed, many studies show that amino acids act heterogeneously and cooperatively within proteins, producing epistasis between amino acids on vastly different scales. At one level, there are local, pairwise epistatic interactions that define direct contacts in the tertiary structure, which likely contribute to native-state stability. But, at another level, there are collectively acting networks of amino acids that mediate central aspects of protein function – binding^15^, catalysis^16^, and allosteric communication^17^. Past work shows that both scales of couplings are represented in the pattern of empirical correlations in multiple sequence alignments (MSAs)^18, 19^, providing different sequence-based methods for understanding protein structure^20^ and function^21^. Thus, a basic requirement for statistical models of protein sequences is to account for both local and collective amino acid epistasis.

A fundamental problem in making such models is the lack of a ground truth for validating all features of the inference process. For example, the local epistasis can can be verified by direct contacts in atomic structures of members of a protein family^1–4^, but a similar benchmark for global collective actions of amino acids is not broadly available. Indeed the inspiration for building statistical models from evolutionary data is, in part, to provide hypotheses for the collective behaviors of amino acids as a route to understanding protein function. How then can we better understand the inference process itself? In this work, we take the approach of using “toy models”^22–24^ in which we (1) specify a pattern of amino acid couplings for a hypothetical protein, (2) generate synthetic sequences that satisfy those constraints, and (3) examine the ability of statistical inference methods to learn these patterns (Fig. 1A-B). This analysis shows that in any practical context, model inference is biased by the limited sampling of sequences. The consequence is that features of different size and strength are unevenly inferred with current methods. These findings are confirmed in a real protein model system in which experimental data allow us to verify both structural contacts and functional amino acid networks. This work clarifies apparent inconsistencies in the current interpretation of coevolution in proteins and opens a path towards new methods for more completely inferring the information content of protein sequences.

**Figure 1.**
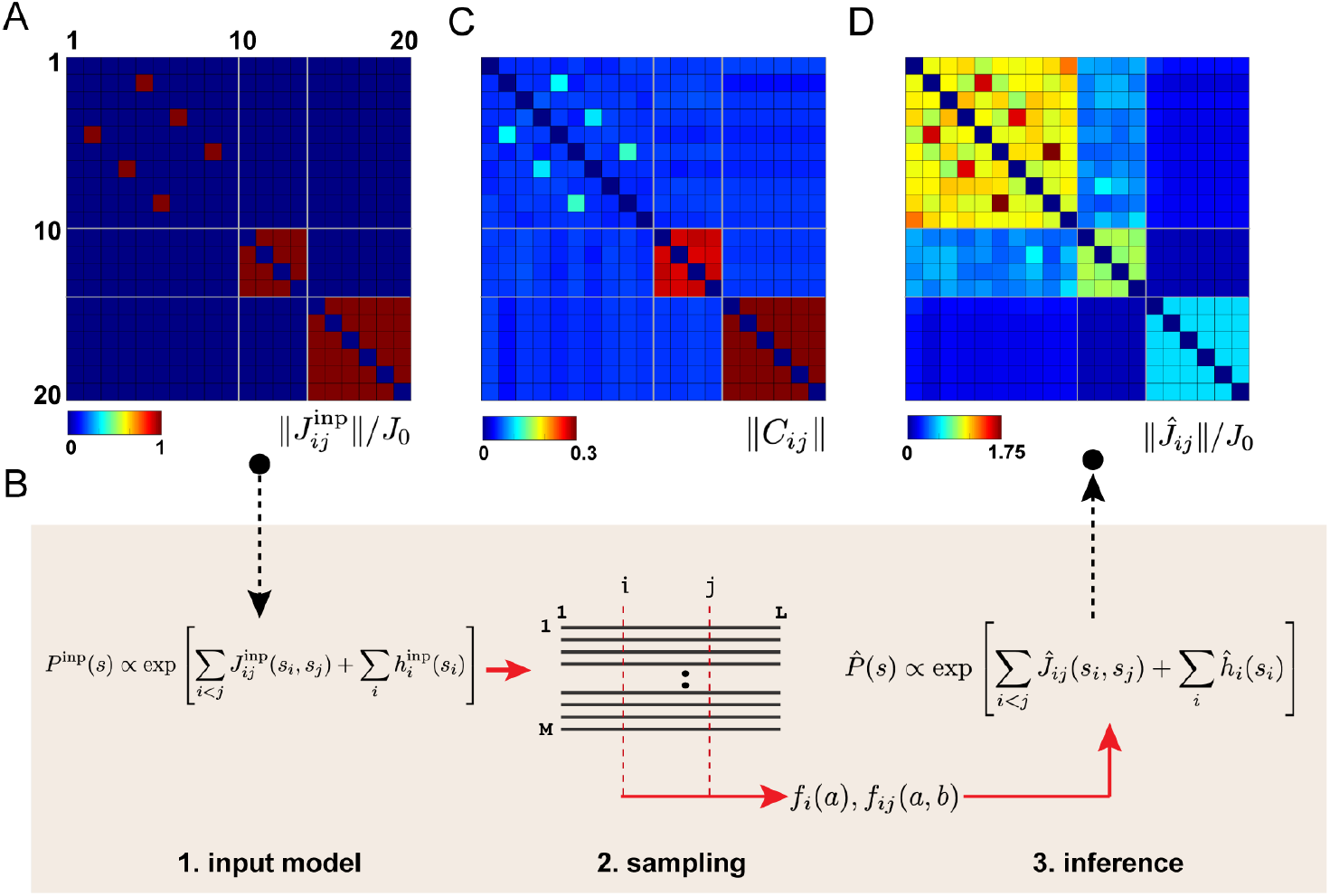
Inference from a toy model of proteins. The model assumes a sequence of length *L* = 20 with *q* = 10 possible amino acids at each position. **A**, the pattern of input couplings between sequence positions. There are three types of features: three isolated pairwise couplings (“contacts”, 2-5, 4-7, and 6-9), a small collective group (“small sector”, all possible couplings within positions 10-13), and a large collective unit (“large sector”, all possible couplings within positions 14-20). All non-zero couplings have same magnitude, see text. **B**, the strategy used in this work, in which we make the input model (step 1), sample *M* sequences from a Boltzmann distribution defined by the input 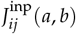 and compute the empirical first and second order statistics *f_i_* (*a*) and *f_ij_*(*a*, *b*) (step 2), and use the DCA approach to infer back the input couplings from the sampled sequences (step 3). **C**, Frobenius norm ‖*C_ij_*‖ of the empirical correlation matrix *C_ij_*(*a*, *b*) = *f_ij_*(*a*, *b*) − *f_i_* (*a*) *f_j_*(*b*) computed from the sampled sequences, showing that the collective groups are most strongly correlated. **D**, the inferred couplings with usual settings in DCA (regularization *λ_J_* = 10^−3^). As described in the text, panels **A** and **D** show normalized couplings 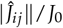.

## Results

### Inference from toy models

A generative statistical model of protein sequences is provided by the Direct Coupling Analysis (DCA)^3, 20^, or more generally a Markov random field. This method starts with a multiple sequence alignment (MSA) of a protein family comprised of *M* sequences by *L* positions, and makes the assumption that each sequence *s* = (*s*_1_,…, *s_L_*) is a sample from a Boltzmann distribution of a Potts model,

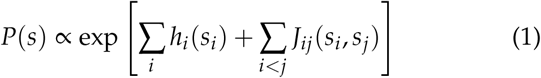

where *h_i_*(*a*) represents the intrinsic propensity of each amino acid *a* to occur at each position *i* (the “fields”), *J_ij_*(*a*, *b*) represents the constraints between amino acids *a*, *b* at pairs of positions *i*, *j* (the “couplings”), and *P*(*s*) is the probability of sequence *s*.

The parameters (*h*, *J*) are inferred by maximum likelihood and are related to the frequencies *f_i_*(*a*) and joint frequencies *f_ij_*(*a*, *b*) of amino acids at positions *i*, *j* by the consistency equations

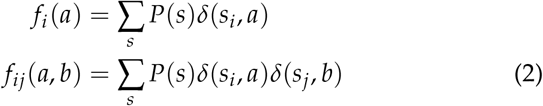

where *δ*(*x*, *y*) = 1 when *x* = *y* and zero otherwise. The probability distribution *P*(*s*) can also be viewed as the maximum entropy model that reproduces the empirical frequencies *f_i_*(*a*) and *f_ij_*(*a*, *b*) of the protein family^3^.

In practice, exact inference of the parameters (*h*, *J*) is computationally intractable because of the number of terms in Eq. (2) is excessively large, but effective approximations exist. In this work, we use pseudo-likelihood maximization (plmDCA)^25^, but the results are consistent with other approximations, including Boltzmann machine learning (bmDCA)^26^. When possible, we make use of exact calculations (Fig. S1).

A critical fact is that in any practical situation, the inference is carried out in the limit of extremely poor sampling. Typically, MSAs contain on the order of 10^3^-10^4^ effective sequences while the number of parameters (*h*, *J*) to be estimated is on the order of 10^5^-10^7^. This undersampling necessitates the use of statistical regularization during the inference process to avoid overfitting. A standard approach is the so-called *L*_2_ regularization, meaning that the log-likelihood function is penalized by a term proportional to the *L*_2_ norm of the parameters. The larger the regularization, the more constrained the parameters. If the fields *h_i_*(*a*) and the couplings *J_ij_*(*a*, *b*) are regularized separately, this changes the consistency equations to

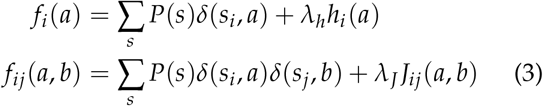

where *λ_h_* and *λ_J_* are the regularization parameters. How does one choose these parameters? Since the inference is unsupervised and cross-validation strategies cannot be applied, the standard approach is to empirically set them by predictive power for various protein properties of interest^9, 25^.

To more formally study the influence of sample size and regularization on the inference process, we made a toy model of a hypothetical protein obeying Eq. (1) with input parameters (*h*^inp^, *J*^inp^), and asked whether these parameters can in fact be inferred from sequences sampled from the model (Fig. 1B). The model comprises *L* = 20 positions and *q* = 10 possible amino acids and has the following characteristics: all fields are set to zero 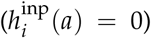, and couplings 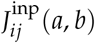 have the pattern shown in Fig. 1A. There are three isolated pair-wise couplings at pairs of positions (2, 5), (4, 7), and (6, 9), a medium-sized interconnected group containing all possible couplings between positions (10-13), and a larger-sized interconnected group containing all possible couplings within positions (14-20). The isolated pairwise couplings mirror the concept of coevolving contacts in protein structures while the interconnected groups of couplings represent the concept of a cooperatively evolving group of positions (sectors). All non-zero couplings have the same strength *J*^inp^ = 2. Note that *J_ij_*(*a*, *b*) is a four-dimensional *L* × *L* × *q* × *q* array, but for presentation, Fig. 1A (and all such panels below) shows the *L* × *L* Frobenius norm 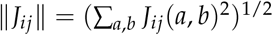 (see methods). We also normalize inferred parameters by the input value, so that perfect inference corresponds to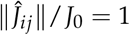 for all non-zero couplings.

We used a Markov Chain Monte Carlo sampling procedure to draw an MSA of *N* = 300 independent sequences from the model (Fig. 1B, step 2), a number that mirrors the undersampling observed in natural protein families. Fig. 1C shows the position by position magnitudes of correlations between amino acids in the sampled sequences. The pattern is heterogeneous, with stronger correlations within the larger interconnected groups of positions. This is because the larger the group, the less likely a position within it is to change over the sampled sequences. In this context, how does DCA work to infer the input couplings *J_ij_*(*a*, *b*) from the empirical statistics? With standard settings for regularization (*λ_J_* = 10^−3^), DCA emphasizes the isolated pairwise couplings while the collective features are hardly discernable relative to noise (Fig. 1D).

### Inference as a function of sample size

Why does DCA selectively emphasize the isolated couplings and under-represent those that make up larger collective features? The answer lies in examining the dependence of the inferred couplings 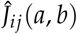 on the degree of sampling in the MSA (Fig. 2A). The data indicate that inferred couplings show three properties as a function of MSA size: (1) they exhibit a “resonance” property where the value of 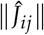 sharply peaks at a characteristic MSA size, (2) they resonate at different characteristic MSA sizes depending on the size of the group they belong to (isolated pairs, small collective, and large collective units), and (3) they only approach their correct values 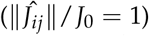 at the limit of very large sampling. At the MSA size chosen in our example (*N* = 300), the isolated couplings dominate the inference, with all collective features many-fold lower in magnitude. However, Fig. 2A also shows that if the MSA contained increasingly more sequences, the outcome would be different; we would suppress the isolated pairwise couplings and instead emphasize the collective features (see Fig. S2 for more details).

**Figure 2.**
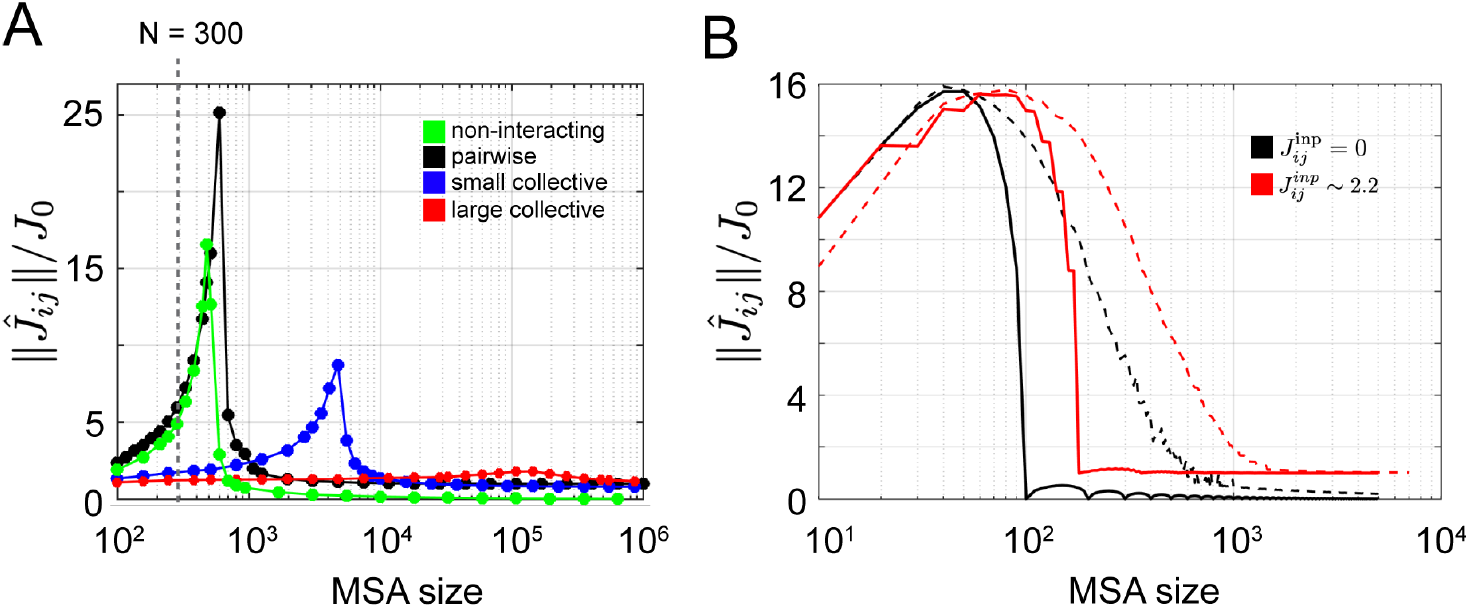
Resonance in inference of sequence features. **A**, normalized magnitude of inferred couplings 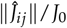 as a function of MSA size, averaged for positions comprising the diffrent sized features in the input model (Fig. 1A). The inference is carried out with weak regularization *J* = 10^−6^). The data show that features in the amino acid sequence display a resonance at characteristic levels of sampling in order of their effective size. Interactions of any size only reach their true values 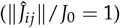 at the limit of infinite sampling. **B**, Inferred couplings for an even simpler model of just *L* = 2 positions and *q* = 10 amino acids either without (black, *J*^inp^ = 0) or with *J*^inp^ ≃ 2.2 (red) input interactions. The traces show cases of deterministic (solid) or random (dashed) sampling of sequences. As described in the text, this model provides a simple mechanistic understanding of the origin of the resonance property.

What is the mechanism of the resonance of inferred couplings as a function of MSA size? To study this, we made an even simpler model of just two positions, each with *q* possible amino acids and with no fields or couplings; that is, with no constraints at all. With infinite sampling, all correlations between amino acids at the two positions must be zero and the inference will return the correct result that all fields a nd couplings are zero. With finite number of sequences, however, the inferred parameters 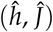 are generally non-zero. For example, consider the situation in which we deterministically draw amino acid pairs uniquely and without repetition to form an MSA of size *N* while keeping amino acid frequencies at both sites uniform. If *N < q*^2^, some amino acid pairs will be observed and the rest (*q*^2^ − *N*) will be absent, requiring inferred couplings in the Potts model to be infinite to account for the absences. The point of regularization is to prevent such an outcome, constraining the difference between the largest and smallest couplings (Δ*J*) to satisfy

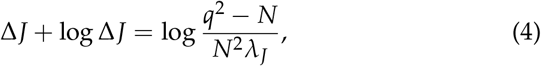

where *λ_J_* is the regularization parameter. It is then easy to show that the magnitude of couplings over all amino acid pairs will be unimodal, with a maximum at the point where the sampling produces the same number of observed and missing pairs (that is, when *N* = *q*^2^/2) (Fig. 2B, solid black curve, and see SI for derivation). The true value of the interaction (*J* = 0) is only reached with complete sampling (*N* ≫ *q*^2^). This shows the basic mechanism of resonance – a sampling-dependent maximization of inferred couplings with a magnitude that is simply set by the strength of regularization (Eq. (4)).

Generalizing to include a non-zero input coupling (red curve in Fig. 2B) has the effect of displacing the resonance curve to the right and shifting the inferred coupling at large sampling to the correct input value (Fig. 2B, compare black and red curves). This makes sense; with stronger coupling, more sampling is generally necessary to draw all possible amino acid states. Thus, as shown in Fig. 2A, the position of a resonance peak is a function of the effective size and magnitude of the input interaction. Pure sampling noise in uncoupled positions resonates at the lowest MSA size, followed in sequence by isolated pairwise couplings and collective features of increasing size. Relaxing the model to use random, rather than deterministic sampling of amino acid pairs just further increases the sampling required for inferring couplings, either without (Fig. 2B, black dashed curve) or with (Fig. 2B, red dashed curve) true interactions.

The implications for sequence-based DCA models of real proteins are clear. Inference for any protein family always occurs at the limit of extreme poor sampling, below the resonance peak for isolated pairwise couplings. In this regime, with usual small regularization to suppress sampling noise, couplings inferred are dominated by the smallest scale features of the information stored in amino acid sequences. Thus the inferred couplings emphasize isolated pairwise couplings between positions that correspond to structural contacts, with couplings within collective groups at or even below the level of pure sampling noise.

The toy model provides another insight into the contact prediction process. A common practice in DCA is to apply an Average Product Correction (APC), which removes a background value from inferred couplings^27^. This approach has been justified by its role in mitigating the effects of phylogenetic bias. However, APC also improves the inference of isolated contacts in our toy model, which includes no notion of phylogeny (Fig. S3). The work here suggests a more general explanation for APC: it works by suppressing the spurious couplings between non-interacting positions that arise due to extreme undersampling. Since in this limit, the noninteracting couplings are comparable in magnitude to the smallest scale of true couplings between positions (Fig. 2), APC helps to separate signal from noise and improve contact prediction in protein structures (Fig. S3).

### Inference as a function of regularization strength

As explained above, the magnitude of inferred couplings in the undersampled regime is basically set by the strength of the regularization parameter *λ_J_* (Eq. (4)). For example, with typical small *λ_J_*, Δ*J* ~ − log *λ_J_*. But how do features of different sizes respond to regularization? To understand this, we carried out model inference for a fixed size MSA (*N* = 300) drawn from the toy model while varying the regularization strength *λ_J_* (Fig. 3A). The data show that for small regularization, the isolated pairwise couplings dominate (black), and collective features (blue and red) are inferred at or below the level of non-interacting pairs (green). As regularization is increased, different features take prominence, until ultimately features are inferred with magnitudes that are in order of their effective size – large collective > small collective > isolated pairs (Fig. 3A). In this strong regularization regime, all inferred couplings decay like 1/*λ_J_* and resemble the empirical correlations *C_ij_*(*a*, *b*) (see SI for details). Remembering that the true input couplings are equal for all features and have normalized value 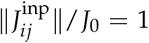, we can conclude that there is no single choice of a regularization parameter that can correctly infer the true pattern of couplings whenever sampling of sequences is limited (compare Fig. 3B-E with Fig. 1A).

**Figure 3.**
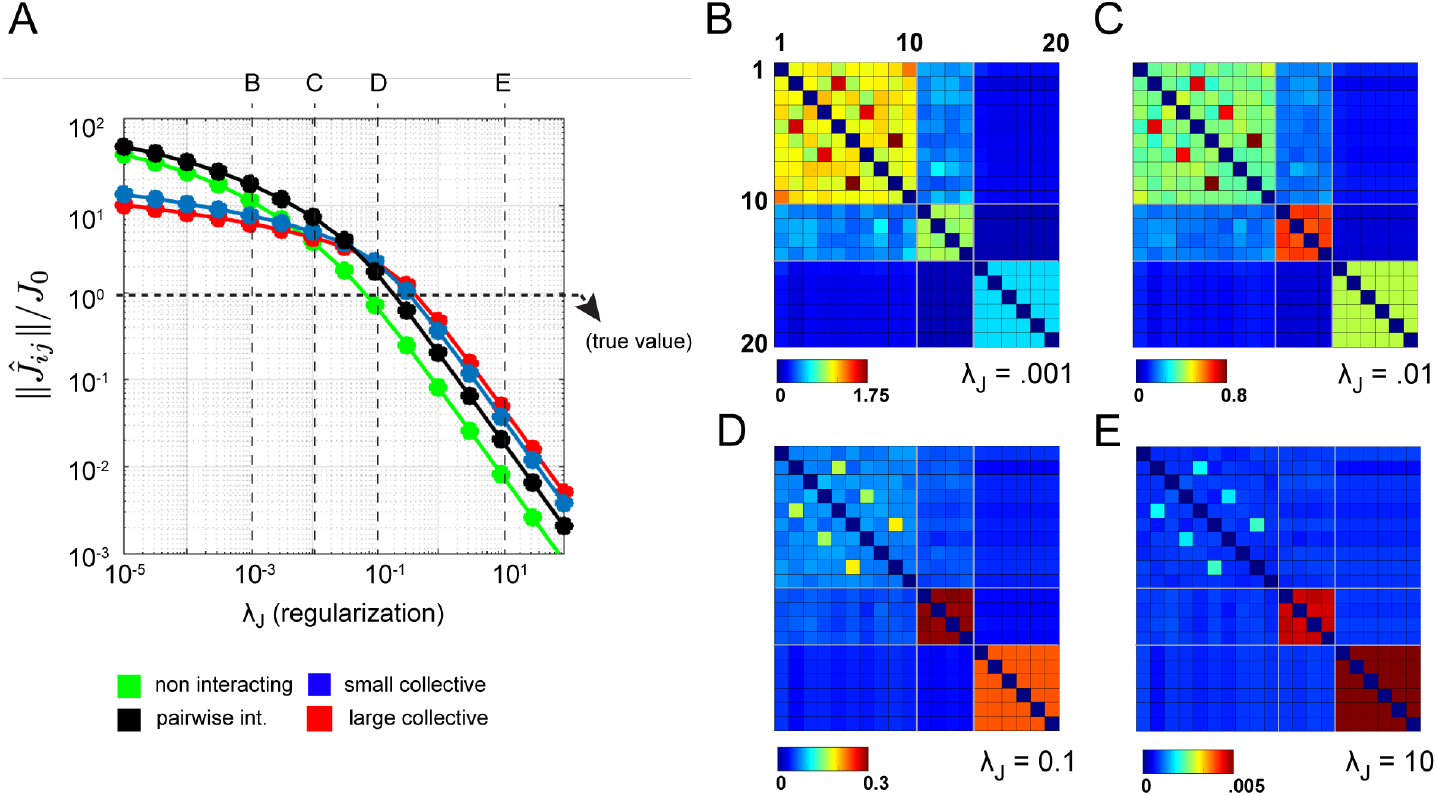
Inference of couplings as a function of the regularization parameter *λ_J_*. **A**, the normalized magnitude of inferred couplings 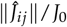 averaged over position pairs comprising isolated pairwise couplings (black), for the small sized collective group (blue), and for the large-sized collective group (red). Inferred couplings for position pairs with zero input couplings are pure undersampling noise and are shown in green. Values of *λ_J_* corresponding to panels B-E are marked, and the true value for non-zero couplings is indicated. **B-E**, for comparison with Fig. 1A, the 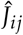 matrix inferred at increasing levels of regularization *λ_J_*. The data show how features of different effective size dominate the inference as regularization is adjusted from small to large values. Note that DCA is traditionally carried out at small regularization strengths *λ_J_* < 10^−2^.

An even simpler model with just two features and two parameters provides an intuitive geometrical illustration of the problem (Fig. 4). This model comprises sequences with *L* = 6 positions and *q* = 2 amino acids with a pattern of input interactions *J*^inp^ shown in Fig. 4A. There is one isolated pairwise coupling between positions 1 and 2 (*J_I_*), and one collective group of couplings between positions 3-6 (*J_C_*) (Fig. 4A), all with the same magnitude 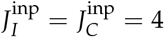. The value of the coupling is chosen simply to be significantly above random fluctuations. This makes the number of parameters to be inferred just two - (*J_I_*, *J_C_*) - enabling us to visualize the inference results on a 2D plane (Fig. 4B). For an undersampled case (here, *N* = 4), the contours of the log-likelihood function being optimized (solid blue contours) show that the inference process has no finite maximum; without regularization, inferred values of couplings *J_I_*, *J_C_* will diverge to infinity. This is consistent with the intuition that couplings must be infinity to account for unobserved amino acid configurations.

**Figure 4.**
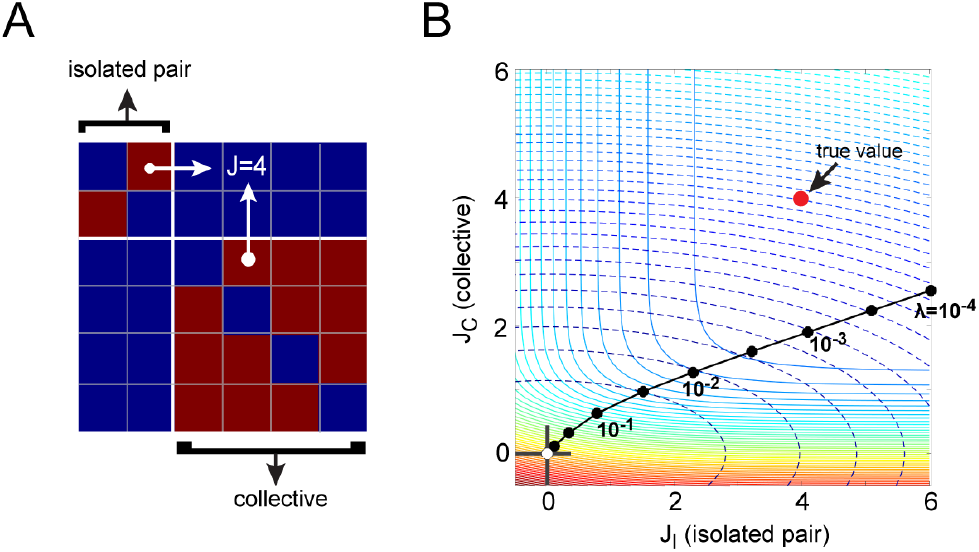
A geometrical explanation of regularized inference. **A**, the input coupling matrix *J*^inp^ for a toy model with *L* = 6 positions and *q* = 2 amino acids and with no fields *h*. The model has two parameters, one representing the isolated pairwise coupling (*J_I_*, positions 1-2) and the other the couplings in the collective set (*J_S_*, positions 3-6). The input values are 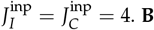. **B**, Inferred values of *J_I_* and *J_C_* from an *N* = 4 undersampled set of sequences for the toy model as a function of regularization *λ_J_*. The solid contours show the landscape of the log-likelihood function being optimized, and the dashed contours show values of 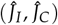 that are consistent with different strengths of regularization.

How does regularization correct this problem? The dashed line contours in Fig. 4B show the curves along which the magnitude of *J_ij_* (that is, 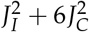) is a constant for various regularization strengths. This defines the solutions to inference with regularization - the points (black filled circles, Fig. 4B) where the solid contours are tangent to the dashed contours. Thus, the inferred solution is set by the regularization used, and there is no regularization at which the inferred solution matches the true solution (*J_I_* = *J_C_* = 4). Also, note that at this level of undersampling, *J_I_* is always larger than *J_C_*. Ananalytical solution relating the regularization parameter *λ_J_* and inferred values of 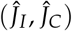 shows how the ratio of these parameters depends on the relative size of the pairwise and collective units, and on the level of sampling (see Supplementary Information).

The bottom line is that with undersampling of data, uniform regularization will always produce solutions that unequally represent the contribution of different sized features and more importantly, that will deviate from the ground truth.

### Application to real problems

These findings have significant impact for model inference in real proteins. The pattern of empirical correlations between pairs of positions in MSAs of protein families reveals a hierarchy of correlation scales, both in terms of magnitude and size of the correlated unit. For example, in an MSA of 1258 members of the AroQ family of chorismate mutase enzymes, a subset of more conserved positions display a pattern of strong interconnected correlations and the remainder of less conserved positions show weaker and more dispersed correlations14 (Fig. 5A). This pattern is reminiscent of Fig. 1C, the correlation matrix resulting from a toy model with features of different effective size. Positions in Fig. 5A are ordered by their susceptibility to regularization (see methods), suggesting that with the extreme undersampling that characterizes all practical MSAs, the inference of couplings in Potts models will inevitably treat these groups unequally. Indeed, just like for the toy model, 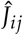 for the AroQ family inferred with standard weak regularization (*λ_J_* = 0.001, Fig. 5B) highlights interactions between the weakly conserved positions while inference with strong regularization (*λ_J_* = 10, Fig. 5C) highlights interactions between the conserved, more collectively evolving positions (Fig. S4). Thus, inference in the context of undersampling selectively represents the information content of protein sequences, with the magnitude of inferred couplings set by the regularization used (see color scale, Fig. 5B-C).

**Figure 5.**
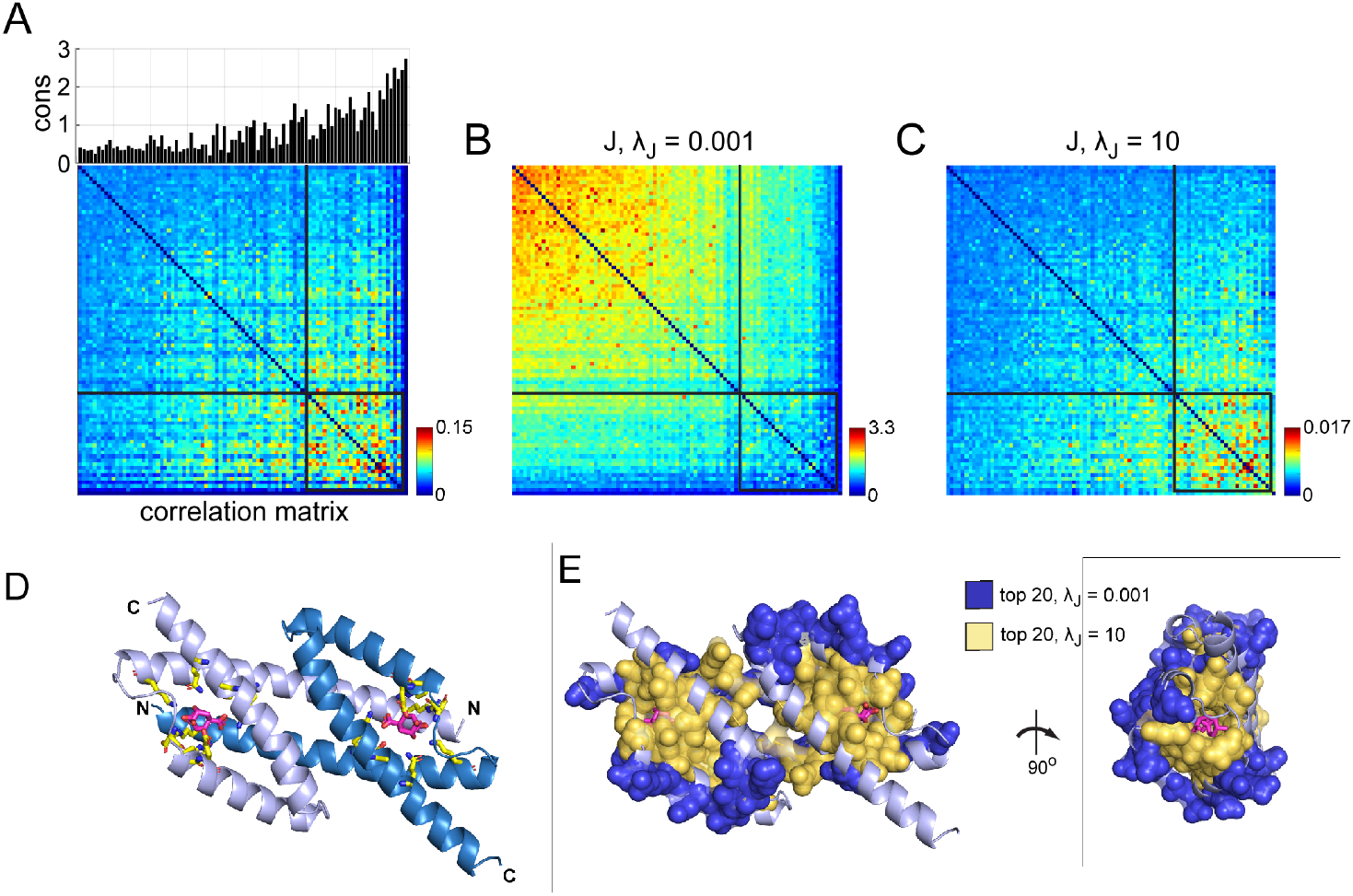
Inference of positional couplings for the AroQ family of chorismate mutase (CM) enzymes. **A**, positional conservation (by Kullback-Leibler relative entropy^21^, bar graph) and the matrix of positional correlations for an MSA of 1258 CM homologs. The positions are ordered by “susceptibility to regularization” (see text and Supplementary Information). **B-C**, the coupling 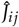 matrix for the CM family inferred with standard small regularization (*λ_J_* = 0.001, panel B) or strong regularization (*λ_J_* = 10, panel C), both ordered as in panel A. **D**, AroQ CMs are dimers with two symmetric active sites formed by elements from both protomers (blue and silver); active site residues are highlighted in yellow stick bonds and a bound substrate analog in magenta. Shown is the structure of the *E. coli* CM domain (EcCM, PDB 1ECM). **E**, spatial organization of positions comprising the top 20 couplings inferred with weak (*λ_J_* = 0.001, blue spheres) or strong (*λ_J_* = 10, orange spheres) regularization.

How do these findings influence our understanding of protein structure and function? AroQ CMs occur in bacteria, archaea, plants, and fungi and catalyze the conversion of the intermediary metabolite chorismate to prephenate, a reaction essential for biosynthesis of the aromatic amino acids tyrosine and phenylalanine. Structurally, these enzymes form a compact domain-swapped dimer of relatively small protomers with two active sites (Fig. 5D). The top terms in 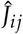 inferred with weak regularization (*λ_J_* = 0.001) correspond to direct contacts between amino acids in the three-dimensional structure (Fig.S5), but are exclusively located within surface-exposed residues (Fig. 5E, blue spheres). In contrast, top couplings inferred with strong regularization (*λ_J_* = 10) switch to represent interactions between buried positions built around the enzyme active site (Fig. 5E, orange spheres). The couplings inferred with strong regulariza-tion include direct tertiary structure contacts, but also comprise indirect, longer-range or substrate-mediated interactions (Fig.S6). The key result is that regularization gradually shifts the pattern of inferred couplings from direct contacts at surface sites to a mixture of direct and indirect interactions within the protein core.

What is the functional meaning of these findings? To comprehensively evaluate this, we carried out a saturation single mutation screen (a “deep mutational scan”) of the AroQ CM domain from *E. coli* (EcCM), following the effect on catalytic activity. This work is enabled by a quantitative select-seq assay for CM activity, reported recently^14^. Briefly, a library comprising all single mutations was made by oligonucleotide-directed NNS-codon mutagenesis, expressed in a CM-deficient *E. coli* host strain (KA12/ pKIMP-UAUC, see Methods), and grown together as a single population under selective conditions. Deep sequencing of the populations before and after selection provides a log relative enrichment score for each mutant relative to wild-type which quantitatively reports the effect on catalytic power^14^. This information is displayed as a heatmap in Fig. 6A - a global survey of mutational effects in EcCM.

**Figure 6.**
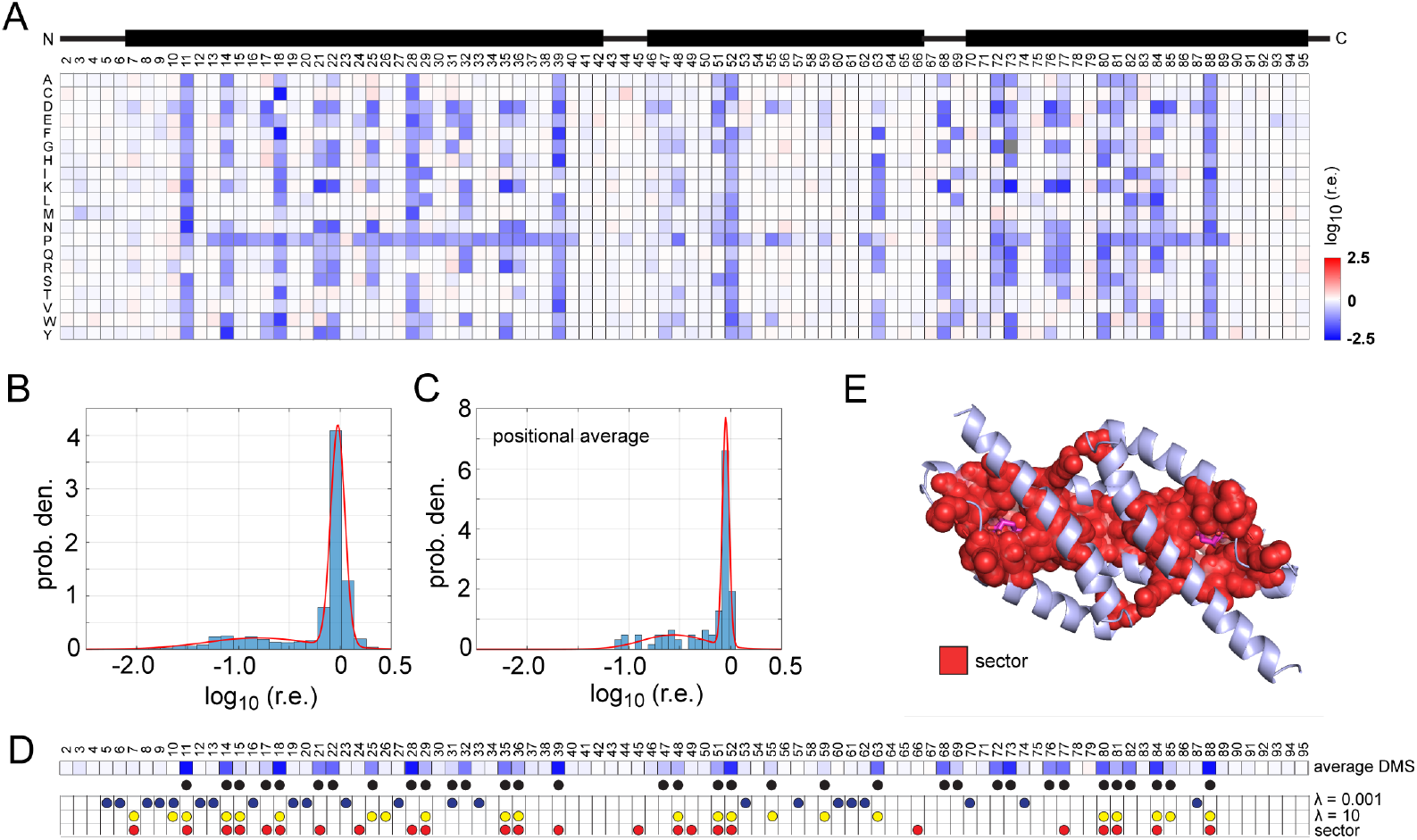
Functional analysis of positions in the *E. coli* CM domain. **A**, A deep mutational scan (DMS), showing the effect of every single mutation on the catalytic power relative to wild-type (see Methods). Blue shades indicate loss-of-function, red indicates gain-of-function, and white is neutral. The cartoon above indicates the secondary structure. **B-C**, the distribution of mutational effects displayed for all amino acid substitutions (B) or for the average effect of mutations at each position (C). The data are fit to a Gaussian mixture model with two components (red curve). **D**, the average effect of mutations is shown as a heatmap and circles below mark the positions comprising the deleterious mode in panel C (black), positions comprising the top 20 couplings inferred with weak (*λ_J_* = 10^−3^, blue) or strong (*λ_J_* = 10, yellow) regularization, and positions comprising the sector as defined by the SCA method (red). **E**, the sector forms a physically contiguous network within the core of the CM enzyme linking the two active sites across the dimer interface.

The distribution of mutational effects is bimodal (Fig. 6B-C), with one mode representing neutral variation and the other representing deleterious effects (black circles, Fig. 6D). The comparison with positions inferred in the top couplings of 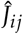 is clear - the top couplings in-ferred with standard weak regularization occur almost exclusively at mutationally tolerant positions, while those inferred with strong regularization occur at functionally important positions (Fig. 6D, *p* = 3 × 10^−8^, Fisher Exact Test) (Fig. S7-S8). Consistent with this, the top couplings inferred with strong regularization significantly o verlap with t he network of conserved, epistatically-coupled, functionally-relevant positions (the sector) defined by the statistical coupling analysis (SCA) method21 (*p* = 1.9 × 10^−7^, Fisher Exact Test) (Fig. 6D-E).

A systematic analysis of the effect of regularization on inference of positional couplings is shown in Fig. 7. The data show that contacts and functional positions are differentially emphasized, with contacts acting like isolated pairwise couplings and functional sites acting like a more epistatic collective unit.

**Figure 7.**
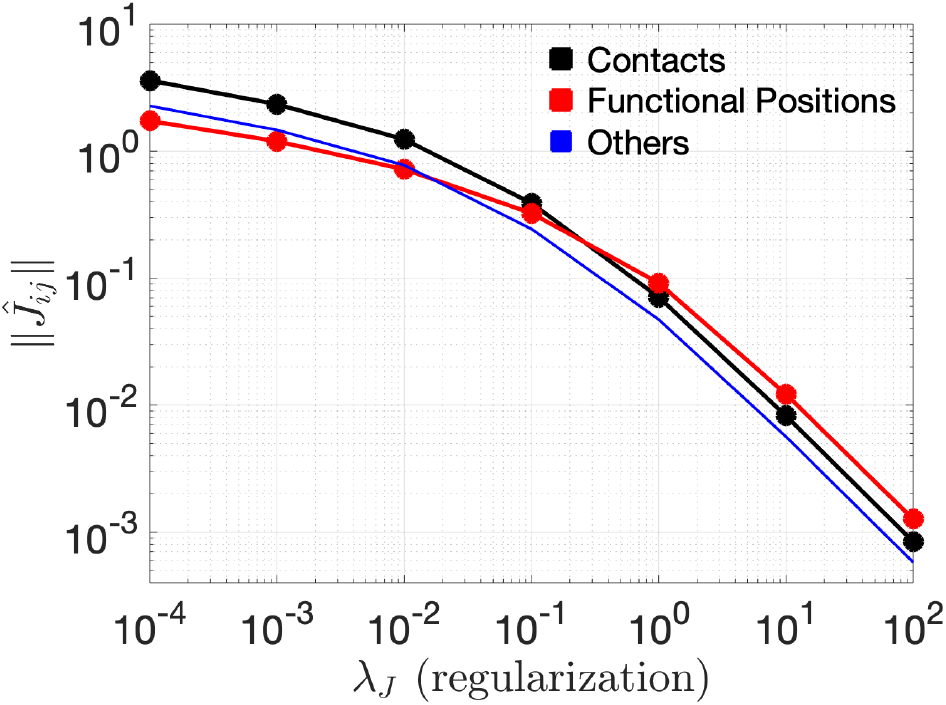
Inferred couplings as a function of regularization *λ_J_* in the CM protein family. The graph shows the magnitude of inferred couplings 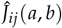 averaged over significant couplings in functional positions as defined in Fig. 6 (red), direct contacts (black), and all other position pairs (light blue). Positions comprising three groups are given in the Suppleentary Information. In analogy with inference for toy models (Fig. 3A), these data show that features of different effective size (here, isolated pairwise contacts and interactions in a collective network) differentially dominate the inference as regularization is adjusted from small to large values.

## Conclusion

The inference of coevolution between amino acids has been valuable, providing new hypotheses for protein mechanisms and global rules for design. One approach is based on Potts models, in which empirical frequencies and correlations of amino acids in a multiple sequence alignment are used to define a probability distribution for the protein family over all sequences^20^. The Potts model has been demonstrated to reveal pairwise tertiary contacts between amino acids^3, 20^, opening the path to sequence-based structure prediction^1, 4^. In this regard, the apparent inability of Potts models to obviously describe collective interactions of amino acids has been puzzling^28^. The collective interactions have been shown to specify native-state foldability^13^, biochemical activities^10, 12, 15, 16, 29, 30^, allosteric communication^17, 31, 32^, and evolvability^33^, defining features of proteins that are essential for their biological function.

The work presented here explains the nature of this problem. With limited sampling of sequences in practically available MSAs, features of different effective size and conservation will be differentially emphasized as a function of MSA size and regularization. With weak regularization, the inference will focus on small-scale, relatively unconserved, functionally-less important local structural interactions, and with strong regularization, inference will emphasize larger-scale, conserved, functionally-essential features. A key point is that there is no single setting of regularization at which all features will be correctly represented. We illustrate this concept using Potts models and one form of regularization (*L*_2_), but the principle of heterogeneous inference of features of different scale is a general one, and depends only on working in the undersampled regime.

The consequence of biased inference is clearly seen in the use of Potts models for protein design. Recent work shows that sequences drawn from a Potts models of chorismate mutate enzymes are indeed true synthetic homologs of the protein family, displaying function both *in vitro* and *in vivo* that recapitulates the activity of the natural counterparts^14^. However, this result required sampling from the model at computational temperatures less than unity, a process that is meant to shift the energy scale to correct for regularization and to enforce under-estimated but functionally essential couplings. This procedure nicely recovers protein function, but does so at the expense of dramatic reduction in sequence diversity of designed proteins compared to natural ones^14^. In light of the work presented here, we can now understand this problem as a non-optimal solution to compensating the unequal inference of features by globally depressing the energy scale.

Can we then “correct” the inference process to more uniformly and accurately represent the biologically relevant patterns of amino acid interactions? Given that practical MSAs will always be grossly undersampled, the main parameter we can control is regularization. But since no single regularization parameter can provide a proper inference of all scales of interactions, it seems clear that what is needed is a strategy for inhomogeneous regularization, where parameters in the Potts model are inferred according to the level of sampling noise that acts on them. If done correctly, such a process should lead to a model that unifies the inference of both local and collective features and enables design of artificial proteins that recapitulate the sequence diversity of natural members of a protein family. The computational toy models introduced here, and the availability of powerful experimental systems such as the chorismate mutases may provide the foundation for this next advancement in sequence based models for proteins.

## Methods

### Toy models and simulated data

Simulated data are generated from input Potts models, with couplings and fields *J*^inp^, *h*^inp^. The inferred couplings and fields are denoted 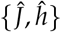. The input model described in Figs. 1–3 involves zero fields 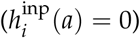 and couplings with non-zero interactions set to equal strength 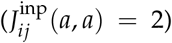. This choice makes the pattern of couplings favorable for *i* and *j* to have identical amino acids, excluding frustration. Sequences are generated from input models through a Markov-Chain Monte Carlo process using the Metropolis-Hastings algorithm. Each sample is obtained after 2 · 10^5^ Monte Carlo iterations starting from independent random sequences. All codes were written in house using MATLAB (Mathworks Inc.) and are available upon request.

### Inference and Gauge

Exact calculations were used for model inference in the small systems described in Figs. 2B and 4. The process involves numerical minimization of the negative log likelihood function, with a regularization term

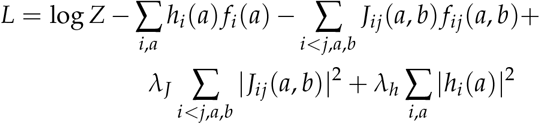

where 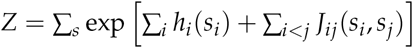 is the partition function, with *s* running over the entire space of sequences.

For all other cases involving larger systems, we used the pseudo-likelihood maximization method plmDCA^25, 34^ for approximate inference, with L2 regularization on both fields (*λ_h_*), and on couplings (*λ_J_*). The value of *λ_h_* is set consistent with past work to be *λ_h_* = 0.01 and the values of *λ_J_* as indicated in the main text.

For sequences of length *L* and number of amino acids *q*, the couplings and correlations comprise four dimensional *L* × *L* × *q* × *q* arrays, and to represent them in as two-dimensional matrices, we take the Frobenius norm over amino acids, defined by

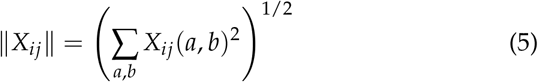

For couplings, this projection is gauge dependent and we implement it in the the zero-sum (or Ising) gauge, such that

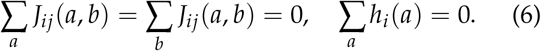

This gauge minimizes the Frobenius norm over all gauges. Note however that the inferred model 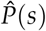 is indepen-dent of the choice of the gauge. For comparison with input values *J*^inp^, the inferred values 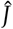 are represented as 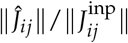 which is by definition 1 when the inference is correct.

### Multiple sequence alignment

Sequences of the AroQ family were acquired by three rounds of PSI-BLAST^35^ using residues 1-95 of EcCM (the chorismate mutase (CM) domain of the *E.coli* CM-prephenate dehrdratase) as the intial query (e-score cut-off 10^−4^). For alignment, we created a position-specific amino acid profile from 3D alignment of four CM atomic structures (PDB IDs 1ECM, 2D8E, 3NVT, and 1YBZ) and iteratively aligned nearest neighbor sequences from the PSI-BLAST using MUSCLE^36^, each time updating the profile. The resulting multiple alignment was subject to minor hand adjustment using standard rules and trimmed sequentially (1) to retain positions present in EcCM, (2) to remove sequences with less than 82 positions, (3) to remove sequences with more than 30% gaps, and to remove excess sequences with more than 90% identity to each other. The final alignment contains 1259 sequences and 96 positions.

### Deep mutation library

A saturation single site mutational library for EcCM was constructed using oligonucleotide-directed NNS codon mutagenesis. To mutate each position, two mutageneic oliognucleotides (one sense, one antisense) were synthesized (IDT) that contain sequences complementary to ~15 base pairs (bp) on either side of the target position and an NNS codon at the target site (N is a mixture of A,T,C,G bases and S is a mixture of G and C). One round of PCR was carried out with either the sense or antisense oligonucleotide and a flanking antisense or sense primer. A second round amplification with first round products and both flanking primers produced the full-length double-stranded product, which was purifed on agarose gel and quantitated using Picogreen (Invitrogen). All first round products were pooled in equimolar ratios, purified, digested with NdeI and XhoI, and ligated into correspondingly digested plasmid pKTCTET-0^37^. For selection, the library was transformed into electrocompetent NEB 10-beta cells (NEB) to yield ¿1000x transformants per gene, cultured overnight in 500 ml LB supplemented with 100 *μ*g/ml ampicillin (Amp), and subject to plasmid purification. The library was diluted to 1 ng/ml to minimize multiple transformation and transformed into the CM-deficient strain KA12 containing the auxilliary plasmid pKIMP-UAUC^38^ to yield ¿1000x transformants per gene. The mixture was then recultured in 500 ml LB containing 100 *μ*g/ml Amp and 30 *μ*g/ml chloramphenicol (Cam) overnight, supplemented with 16% glycerol, and frozen at −80◦C.

### Chorismate mutase selection assay

The selection assay followed a recently reported protocol^37^. Briefly, glycerol stocks of KA12/pKIMP-UAUC carrying the saturation mutation library in pKTCTET-0 were cultured overnight at 30◦C in LB supplemented with 100 *μ*g/ml Amp and 30 *μ*g/ml Cam. The culture was diluted to *OD*_600_ of 0.045 in M9c minimal medium37 supplemented with 100 *μ*g/ml Amp, 30 *μ*g/ml Cam, and 20 *μ*g/ml each of L-phenylalanine (F) and L-tyrosine (M9cFY, non-selective conditions), grown at 30°C to *OD*_600_ ∼ 0.2, and washed in M9c (no FY). An aliquot of the washed culture was used to innoculate 2 ml LB with 100 *μ*g/ml Amp, and grown overnight at 37°C and harvested for plasmid purification (the pre-seleted, or input sample). For selection, another aliquot of the washed culture was diluted to a calculated starting *OD*_600_ = 10^−4^ into 500 ml M9c supplemented with 100 *μ*g/ml Amp, 30 *μ*g/ml Cam, 3 ng/ml doxycycline (to induce CM gene expression from the *P_tet_* promoter) and grown at 30°C for 24h to a final *OD*_600_ < 0.1. Fifty ml of the culture was harvested, resuspended in 2 ml LB with 100 *μ*g/ml Amp, grown overnight at 37°C, and harvested for plasmid purification (the seelcted sample).

Input and selected samples were amplified using two rounds of PCR with KOD polymerase (EMD Millpore) to add adapters and indicies for Illumina sequencing. Amplification in the first round included 6-9 random bases to aid initial focusing and part of the i5 or i7 Illumina adapters. The remaining adapter sequenes and TruSeq indicies were addred in the second round. PCR was limited to 16 cycles and included high initial tempate concentration to minimize amplification bias. Final products were gel purified (Zymo Research), quantified by Qubit (ThermoFisher), and sequenced on an Illumina MiSeq system with a paired-end 250 cycle kit. Paired-end reads were joined using FLASH, trimmed to the NdeI and XhoI cloning sites and translated. Only exact matches to library variants were counted. Relative enrichments (r.e.) were calculated according to the equation 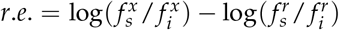 where 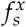 and 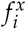 represent the frequencies of each allele *x* in either selected (*s*) or input *i* pools and 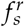 and 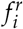 represent those values for EcCM, the wild-type reference.

## Supporting information

Kleeorin_SI

## Acknowledgments

We thank M. Weigt, R. Monasson, S. Cocco, F. Zamponi, A.F. Bitbol, Y. Meir, N.S. Wingreen and members of the Ranganathan and Rivoire laboratories for discussions. This work was supported by grant FRM AJE20160635870 (O.R.), grant ANR 17-CE30-0021-02 (O.R.), NIH grant RO1GM12345 (R.R.), a Data Science Discovery Award from the University of Chicago (R.R.) and a collaboration grant from the France-Chicago Center (R.R. and O.R.).

## Notes

### Competing Interest Statement

The authors have declared no competing interest.

