## Supplementary material for "Undersampling and the inference of coevolution in proteins": Kleeorin_SI

### Comparison between methods of inference

As noted in the main text, the inference of Potts models is computationally intractable for all but the smallest of systems. The calculations involve the estimation of the marginals $\hat{f}_i(a), \hat{f}_{ij}(a,b)$ as a function of the model parameters $\{\hat{J}, \hat{h}\}$ and an exact estimation requires a sum over the space of all possible sequences (see Eq. (3) of the main text). Several approximations have been proposed to address this computational problem (1), including the plmDCA method and Botlzmann machine learning (bmDCA, (2)). In the main text, we use an exact calculation for Fig. 2B and Fig. 4 and the plmDCA method otherwise. Fig. S1 shows comparisons of inference with these various approaches, demonstrating robustness of the claims in this work to the method of inference.

### Lower bound for minimal sampling

The effects that we describe in the main text arise from under-sampling, such that certain combinations of pairs of amino acids are not represented in the data. The typical number of samples needed to overcome this problem depends on the specifics of the generating models; for example, models with stronger constraints, meaning larger couplings and/or larger collective units will require a larger number of samples. This phenomenon lies at the heart of the heterogeneity in sampling noise experienced by features of different effective size in practical multiple sequence alignments. To obtain an analytical expression for the lower bound on sampling, we consider here the least constrained model, which is a null model with no fields or couplings at all, and estimate the mean number of samples needed to observe all possible pairs of amino acids at least once, as a function of the number $L$ of positions and the number $q$ of possible amino acids.

The numerical results (Fig. S2) suggest a scaling with $q$ as $q^2$ and a scaling with $L$ as $\ln L$. These scaling relationships can be understood by a rough calculation that treats the combinations of amino acids independently. Starting with just one pair of positions ($L = 2$), a particular combination $(a, b)$ of amino acids has probability $1/q^2$ of occurring in any particular sample, and the probability that it is not observed in $N$ samples is therefore $(1 - q^{-2})^N \simeq \exp(-N/q^2)$. Treating all combinations of amino acids independently, the probability that one of the $q^2$ combinations is not observed is then $P(N) \simeq 1-(1-\exp(-n/q^2))^{q^2} \simeq q^2 \exp(-N/q^2)$. The necessary number of samples $N_{\min}$ needed to observed all combinations of amino acids thus scales with $q$ as $q^2$.

Extending the argument to $L > 2$ positions under the same simplifying assumption that combinations of amino acids can be independently, the total number of combinations becomes $q^2L(L-1)/2$ and $P(N) \simeq q^2L(L-1)/2\exp(-N/q^2) \simeq \exp(2\ln q + 2\ln L - N/q^2)$, from which it follows that the required number of sequences $N_{\min}$ scales with $L$ as $\ln L/L_0$.

For a length $L = 100$ and $q = 20$ amino acids, this corresponds to $N_{\min} \simeq 10^4$ sequences. This is, however, only a lower bound that assumes a model with no constraints. Constraints can increase this number by many orders of magnitudes. For instance, with the model of Fig. 2A, where $L = 20$ ad $q = 10$, the unconstrained model requires $N_{\min} \simeq 10^3$, which is beyond the peak corresponding to isolated couplings but before the peak corresponding to couplings involved in collective units.

Methods such as flavor reduction (3), which effectively decrease $q$, or pseudo-counting (4), which effectively increase $N$, can alleviate under-sampling but come with biases of their own. Note also that pseudo-counts are formally equivalent to L2 regularization in the context of Gaussian models (5).

### Deterministic minimal model for the resonance in the undersampled regime

The "resonance" observed in Fig. 2B arises in an undersampling regime where the log-likelihood has no extremum, and where the results are entirely determined by the regularization. This is illustrated by Eq. (4) of the main text where we consider a minimal model of just $L = 2$ positions with $q$ possible amino acids and no constraint. To focus on the desired effect and isolate it from the contribution of sampling stochasticity, we analyze sets of sequences that have a uniform number of each of the $q$

amino acids, $f_i(a) = f_i(b)$ for any $a, b$ at each position $i$. We further assume that each combination $(a, b)$ is either present or observed in a single sequence for each pair $i, j$. This defines a "deterministic" sampling procedure.

In this scenario, the inferred couplings $\hat{J}_{ij}(a, b)$ can only take two values, depending on whether the combination of $(a, b)$ is observed or not at $(i, j)$. Using Eq. (3) in the main text, this difference $\Delta J$ is given by

$$\frac{1}{N} = \frac{e^{\Delta J}}{Ne^{\Delta J} + q^2 - N} + \lambda_J \Delta J. \qquad \text{[S1]}$$

Assuming that $\lambda_J$ is small, an expansion for large $\Delta J$ leads to Eq. (4) in the main text, .

In the Ising gauge, the inferred couplings for the occurring and missing combinations of amino acids are, respectively, $(1 - N/q^2)\Delta J$ and $-N\Delta J/q^2$. The Frobenius norm of the couplings is therefore maximal when there is the same number of missing and occurring pairs, at $N = q^2/2$. The exact position of the "resonance" peak position is gauge and representation dependent, but the mechanism that leads to a non-monotonic dependence in sampling size is general.

The dependence of the resonant peak on the input coupling $J^{\text{input}}$ can be studied by adding a strong ferromagnetic coupling between the two positions of the model. Most generated sequences then involve the beneficial ferromagnetic combinations of amino acids, which increases the number of samples needed to observe all combinations, and therefore shifts the position of the peak to larger values of sampling size, as seen in Fig. 2B (red).

### Strong regularization limit

In the strong regularization limit $\lambda_J \to \infty$ where $\hat{J}_{ij} \to 0$, we have $\hat{C}_{ij} \simeq \hat{J}_{ij}$ where $\hat{C}_{ij}(a, b) = \hat{f}_{ij}(a, b) - \hat{f}_i(a)\hat{f}_j(b)$ is the correlation obtained from the inferred model $\hat{P}(s)$. Using $f_{ij} = \hat{f}_{ij} + \lambda_J \hat{J}_{ij}$ from Eq. (3), we have therefore

$$C_{ij}(a, b) = \hat{J}_{ij}(a, b) + \hat{f}_i(a)\hat{f}_j(b) - f_i(a)f_j(b) + 2\lambda_J \hat{J}_{ij}(a, b). \qquad \text{[S2]}$$

and

$$\hat{J}_{ij}(a, b) = [C_{ij}(a, b) + f_i(a)f_j(b) - \hat{f}_i(a)\hat{f}_j(b)]/(2\lambda_J + 1) \qquad \text{[S3]}$$

In the strong regularization limit, the inferred couplings are thus proportional to the correlations, up to the addition of a rank-two correction. This correction is controlled by $\lambda_h$, the regularization parameter for the fields, and is negligible whenever the model reproduces the first order statistics, i.e., $\hat{f}_i(a) = f_i(a)$ for all $i, a$.

### The two-parameter minimal model

In Fig. 4B, we analyze a minimal model with $L = 6$ positions, $q = 2$ amino acids and a pattern of couplings indicated in Fig. 4A. This model has only two parameters. One parameter, $J_I$, is the coupling of the isolated pair, and the other parameter $J_C$ is the coupling of pairs in the collective unit. At variance with the model of Figs. 1-3, we ignore all other possible couplings and impose the couplings within the collective units to be identical. Under these assumptions, the model consists in two independent sub-models, each with a single parameter.

We generate a very small data-set of $N = 4$ sequences with input couplings $J_I^{\text{inp}} = J_C^{\text{inp}} = 4$. We typically obtain a data-set where every sequence has the maximal fitness of $F_{\max} = J_I + 6J_C$. We also assume that amino acids are uniformly represented at each position to focus on the inference of the couplings.

To estimate $\hat{J}_I$ and $\hat{J}_C$, consider more generally inferring a common coupling $\hat{J}_n$ between $n$ positions based on the knowledge of a mean fitness given by $F_n = \binom{n}{2} J_n$, ($J_I \equiv J_2$ and $J_C \equiv J_4$). The partition function in the low-temperature limit is $Z_2 = 2(e^{J_2}+1)$ and $Z_n = 2e^{\binom{n}{2}J_n}(1+ne^{-(n-1)J_n})$ for $n > 2$ and the regularized log-likelihood function is $L_{\lambda_J}/N = \ln(1/Z_n) + \binom{n}{2}J_n - \binom{n}{2}\lambda J_n^2$. Differentiating with respect to $J_n$ and keeping again only leading term in $J_n$, we thus obtain $1/\lambda_J = \hat{J}_2 e^{\hat{J}_2}$ and $1/\lambda_J = \hat{J}_n e^{(n-1)\hat{J}_n}$ for $n > 2$. Applied to our minimal model, this gives

$$1/\lambda_J = 2\hat{J}_I e^{\hat{J}_I} = \hat{J}_C e^{3\hat{J}_C} \qquad \text{[S4]}$$

which directly indicates the presence of a bias between $\hat{J}_I$ and $\hat{J}_C$. The ratio $\hat{J}_I/\hat{J}_C$ can be roughly estimated to be $\simeq 3$ in the limit where $\lambda_J$ goes to zero. Fig. 2B indeed indicates an asymptotically linear relationship with $\hat{J}_I/\hat{J}_C \simeq 2.55$.

### Average product correction

An average product correction (APC) is routinely used to predict contacts from the inferred couplings $\hat{J}_{ij}$ (6), where pairs of positions are not scored by the Frobenius norm $\|\hat{J}_{ij}\|$ but by

$$\|\hat{J}_{ij}\|_{\text{APC}} = \|\hat{J}_{ij}\| - \frac{\sum_k \|\hat{J}_{ik}\| \sum_k \|\hat{J}_{kj}\|}{\sum_{kl} \|\hat{J}_{kl}\|} \qquad \text{[S5]}$$

The correction is aimed at removing a background value shared by positions $i, j$. The comparison of Fig. S3 with Fig. 3 shows that APC is indeed effective to enhance the identification of isolated coupled pairs in the low-regularization limit. On the other hand, APC may not be necessarily useful for identifying large collective units: for instance, Figs. S3B,C show that APC seems to highlight smaller scale patterns at the expense of larger scale patterns. As described in the main text, our work suggests that APC mainly works by removing spurious signals that arise by the smallest scale features in an alignment, the non-interacting positions. Since isolated positions scale closest to this random signal (Fig. 2A), the APC is primarily effective at contacts prediction.

### Inference from real data

Figs. 5 and 7 present results obtained by inferring a Potts model from a multiple sequence alignment of chorismate mutases (CMs) previously described in (7), using a standard sequence weighting parameter $\theta = 0.8$ to reduce proximal phylogenetic effects. The inference is performed with plmDCA for different values of $\lambda_J$ while keeping $\lambda_h = 0.01$ fixed. This value, though important, does not influence the results significantly as long as it is kept sufficiently low.

Coupling matrices with positions ordered along the primary sequences are represented in the top row of Fig. S4. In Fig. 5B-C and the bottom row of Fig. S4, the positions are re-ordered to emphasize the differences arising from different choices of $\lambda_J$. The new order is based on a susceptibility to regularization measure, defined by

$$\chi_i = \frac{\sum_j \|\hat{J}_{ij}^{(\lambda_J=10^2)}\|}{\sum_j \|\hat{J}_{ij}^{(\lambda_J=10^{-7})}\|} \quad \text{[S6]}$$

where $\hat{J}_{ij}^{(\lambda_J=x)}$ indicates the coupling inferred with $\lambda_J = x$ and where we compare here two extreme values of $\lambda_J$. The result in the ordered case shows how the protein positions seemingly decompose into two parts that are analogous to Fig. 3, where the cooperative unit and the isolated pairwise couplings switch their relative importance. Note that if pairwise coupling represented the only signal in the data, we would expect a different picture: as regularization increases, the spurious signal would decrease and separate from true signals but the largest couplings would remain the same as when inferred with low regularization. The dramatic switching of couplings between different groups of positions strongly argues for the presence of a heterogeneity of scales in real proteins.

### Interpretation of top couplings

The top $L/2$ couplings inferred at low regularization typically represent contacts in three-dimensional structures (4). We examined the top 20 pairs $(i, j)$ with largest $\|\hat{J}_{ij}\|$ for the CM MSA, applying standard practices for contact prediction - average product correction (see section F, above) and excluding pairs that are less than four positions apart along the linear sequence ($|i - j| > 3$). Figs.S5 and S6 represent the positions that contribute to the top 20 pairs with either weak ($\lambda_J = 0.001$) regularization or strong ($\lambda_J = 10$) regularization. The data show that inference with weak regularization nearly exclusively identifies direct contacts in the tertiary structure of *E.coli* CM (EcCM, PDB 1ecm) but that inference with strong regularization identifies a mixture of direct and indirect or substrate-mediated interactions. Here a contact is defined as two residues with at least one pair of atoms approaching within the sum of their van der Waals radii plus 20 % in the crystal structure of EcCM.

To examine the relationship of couplings to function, we define "experimentally significant" couplings as those comprising the 34 positions comprising the deleterious mode in Fig. 6C, defined by fitting the data to a Gaussian mixture model. This allows us to examine how top pairs found with different levels of regularization are related to CM function. The data show that top pairs obtained in the low-regularization limit correspond to contacts and not to experimentally significant pairs (Fig. S7A; see below for results with an average product correction that leads to a greater number of top pairs that are contacts, as well as a few in the cooperative network). In contrast, 74% of top pairs obtained in the high-regularization limit are experimentally significant pairs, and only 62% are contacts (Fig. S7C). These contacts are distinct from those obtained with weak regularization and overlap with sector pairs (Fig. 5F).

Repeating the analysis of Fig. S7 with the APC, we verify that contact prediction is significantly improved (Fig. S8). In particular, at low regularization, 100% of the top $L/2$ pairs are contacts, as typically found in other protein families (Fig. S8A). In contrast, at high regularization, contact prediction is improved but the inference of experimentally significant pairs drops from 74% to 56% (Fig. S8C). Note, however, that with or without APC, 88% of top pairs can be interpreted as contacts or experimentally significant pairs, suggesting that APC is not just "cleaning noise" but shifting the emphasis from one type of significant pairs to another, namely from pairs in cooperative clusters to pairs in isolated contacts.

In Fig. 7, we define 32 "statistically significant contacts" as top pairs obtained in the low-regularization limit with APC that are in the contact map, but are not part of the functional network. Also, we define 13 "statistically significant experimental couplings" as top pairs obtained with strong regularization without APC that are in the functional network, but are not contacts.

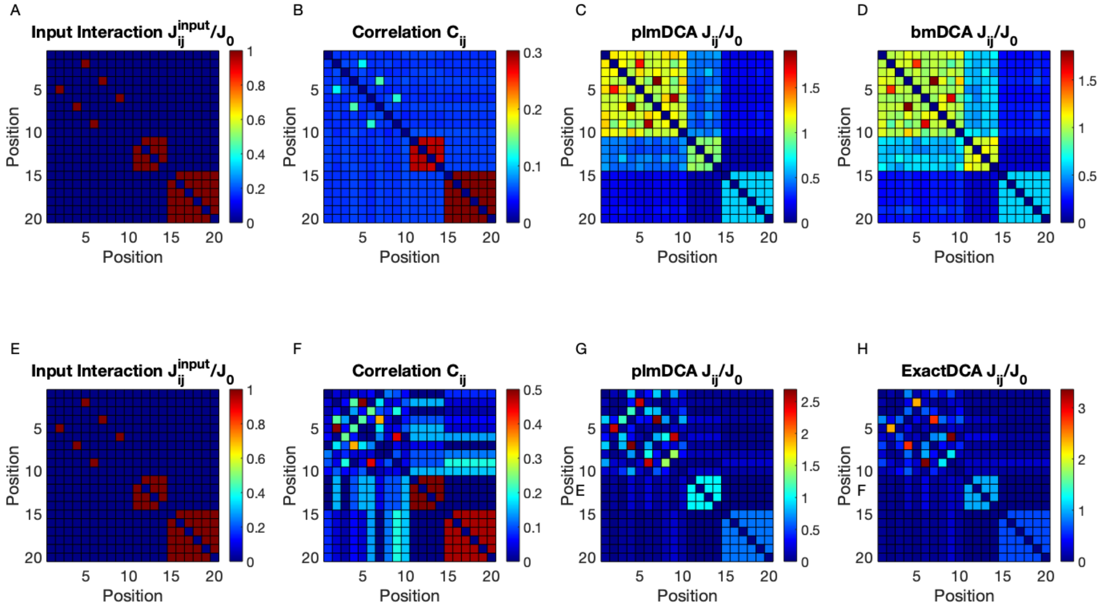


**Fig. S1.** Comparing inference methods – For the same system as in Fig. 1, **(A)** the input positional interaction $J_{ij}^{\mathrm{input}}$ used in generating the data, **(B)** the positional correlation $C_{ij}$ and **(C)** the plmDCA inferred positional couplings $\hat{J}_{ij}$ for $\lambda_J = 10^{-3}$. **(D)** same as (C) but using bmDCA with $5.10^4$ iterations. For the same system, but with $q = 2$ amino acids and only $N = 20$ sequences, **(E)** the input positional interaction $J_{ij}^{\mathrm{input}}$ used in generating the data, **(F)** the positional correlation $C_{ij}$ and **(G)** the plmDCA inferred positional couplings $\hat{J}_{ij}$ for $\lambda = 10^{-3}$. **(H)** is the same as (C) inferred using an exact calculation with same regularization parameter.

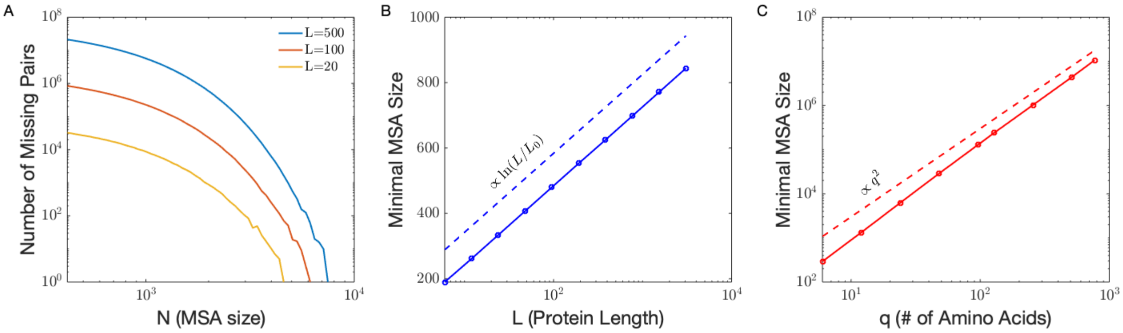


**Fig. S2.** Critical undersampling for unconstrained models - **(A)** Number of missing pairs as a function of MSA size for $q = 21$ and various sequence lengths $L$. We can see that it takes on the order of $10^4$ pairs to minimally sample a totally random MSA. **(B)** Average minimal MSA size needed to avoid the critical undersampling regime in unconstrained models as a function of the sequence length $L$, for $q = 5$. We observe a scaling in $\ln(L/L_0)$. **(C)** Average minimal MSA size to avoid the critical undersampling regime in unconstrained models as a function of the number $q$ of possible amino acids, for $L = 10$. We observe a scaling in $q^2$.

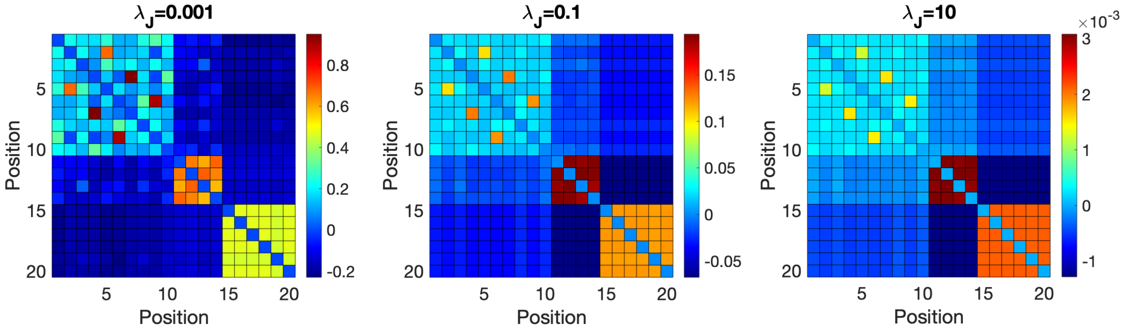


**Fig. S3.** Counterpart of Fig. 3B,D,E using the APC score $\|\hat{J}_{ij}\|_{\mathrm{APC}}$ defined in $\mathrm{Eq.}$ (S5) instead of the Frobenius norm $\|\hat{J}_{ij}\|$.

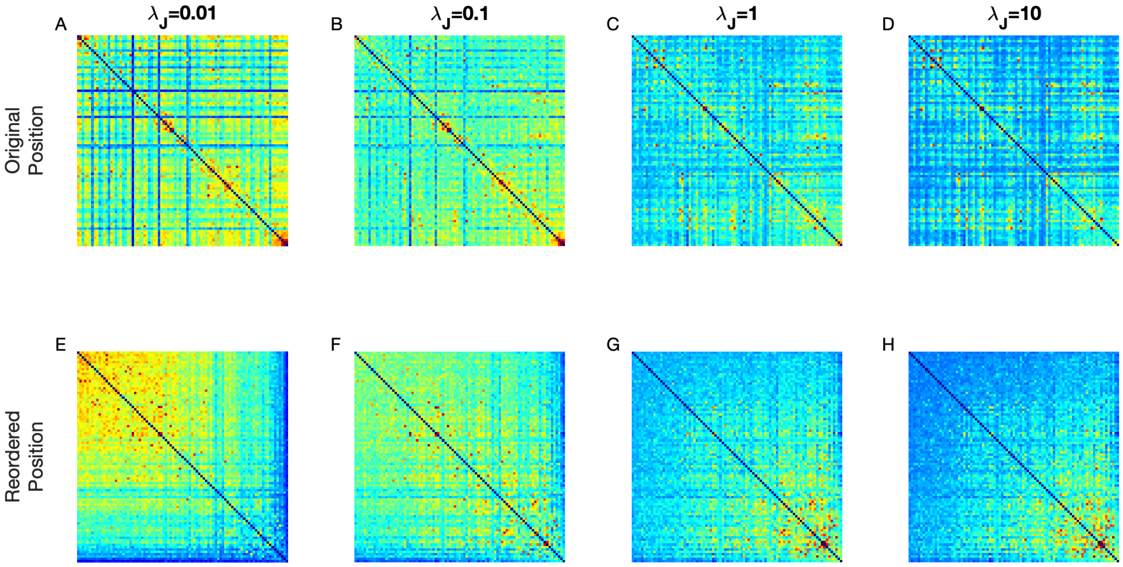


**Fig. S4.** Coupling matrices $\hat{J}_{ij}$ inferred from real data for increasing values of the regularization parameter $\lambda_J$ – In the top row **(A-D)**, positions are ordered as in the linear sequence while in the bottom row **(E-H)** they are ordered by the value of $\chi_i$, defined in Eq. (S6) to represent the sensitivity to change in the regularization parameter $\lambda_J$. As regularization increases, the bottom row shows a tendency analogous to Fig. 3, where the various components of the toy model switch their relative importance

**Top 20 couplings, λ = 0.001**

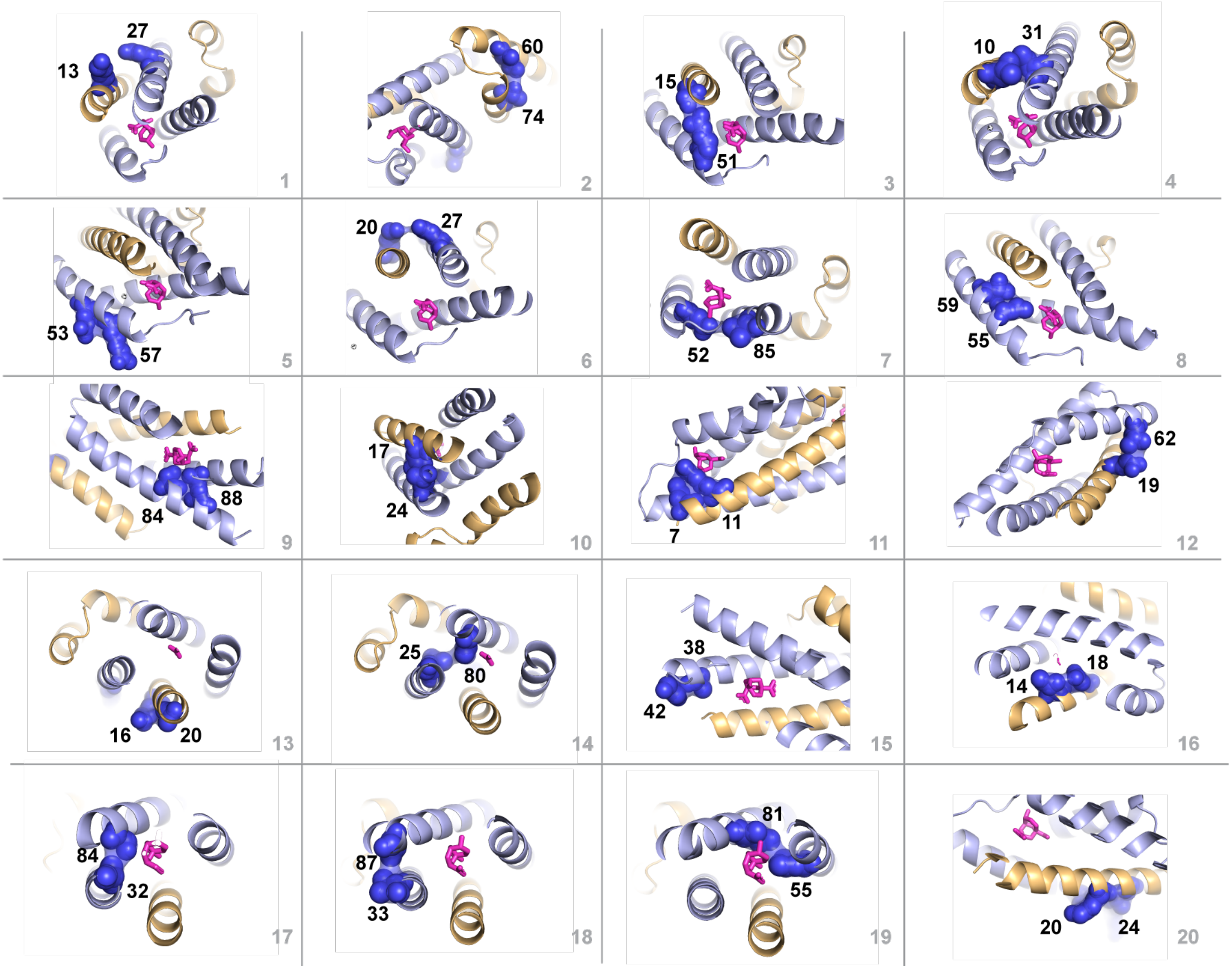


**Fig. S5.** Position pairs comprising the top 20 couplings in $\hat{J}_{ij}$ inferred with weak regularization ($\lambda_J = 0.001$), and with standard practices for contact prediction - average product correction and exclusion of couplings with sequence separation less than four. Nearly all pairs represent direct tertiary structure contacts.

**Top 20 couplings, λ = 10**

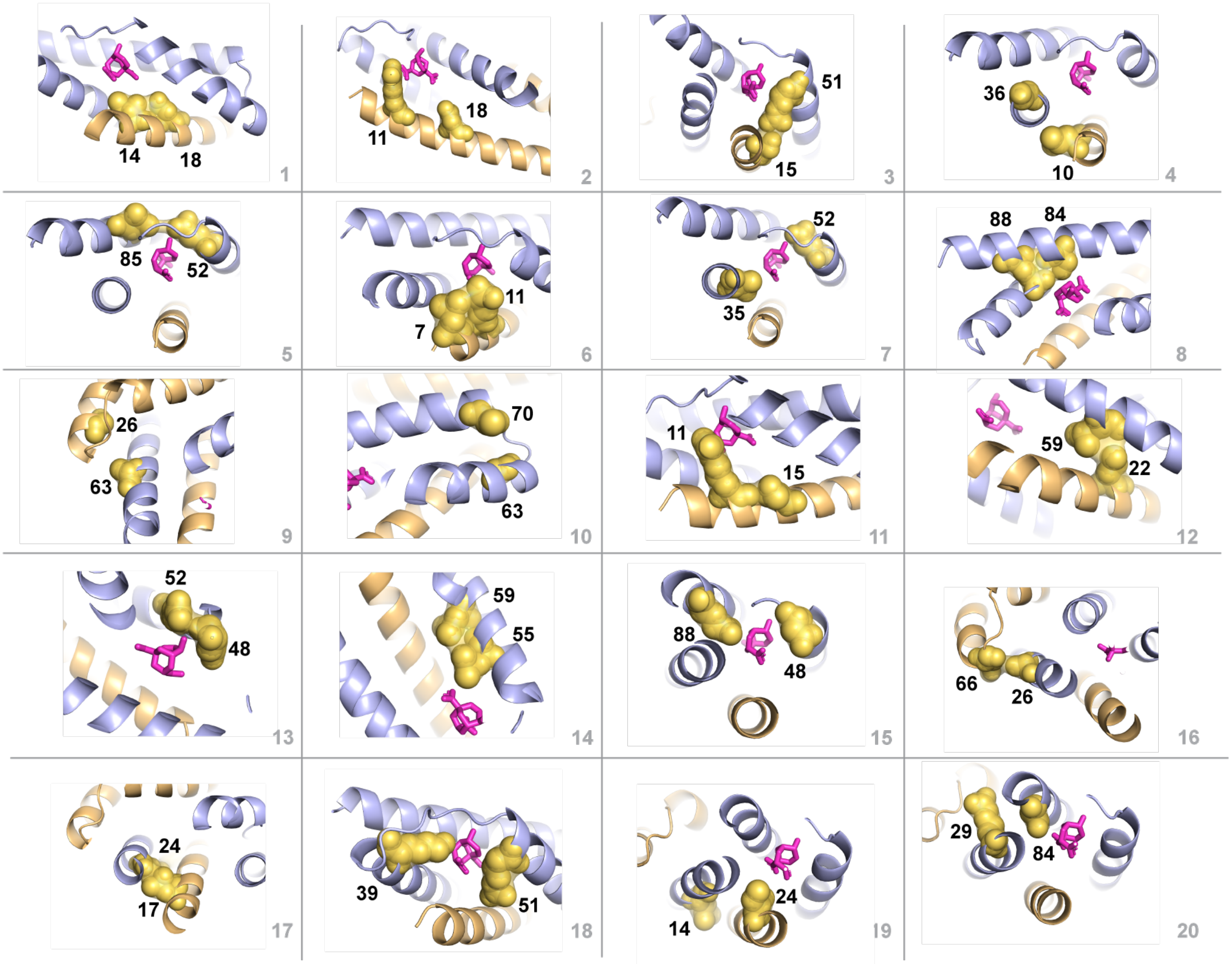


**Fig. S6.** Position pairs comprising the top 20 couplings in $\hat{J}_{ij}$ inferred with strong regularization ($\lambda_J = 10$), and with standard practices for contact prediction - average product correction and exclusion of couplings with sequence separation less than four. Some pairs represent direct tertiary structure contacts, but others show indirect or substrate-mediated interactions.

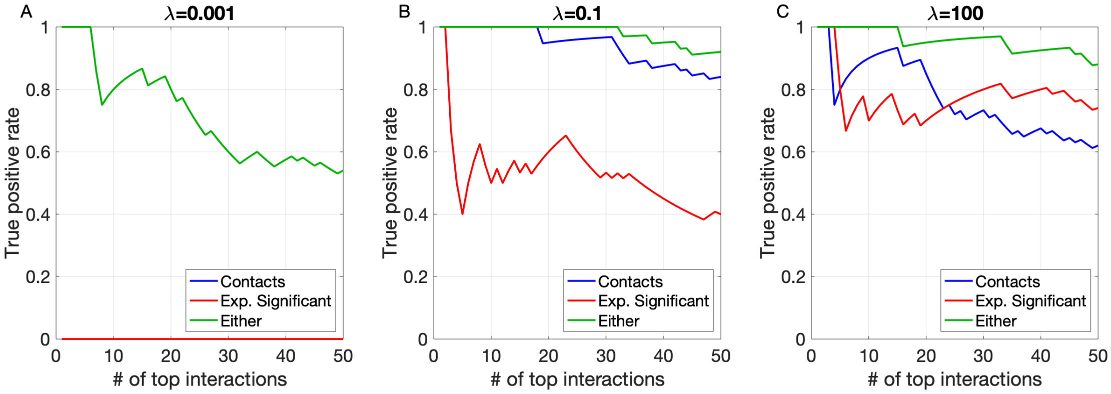


**Fig. S7.** Statistics over top coupled pairs, ranked by $\|\hat{J}_{ij}\|$, the Frobenius norm of the couplings $J_{ij}(a,b)$ – $\mathbf{(A,B,C)}$ For three values of the regularization parameter $\lambda_J$, fraction of top $\|\hat{J}_{ij}\|$ pairs that are contact position pairs (in blue), experimentally sensitive position pairs (in red) or in either one of these groups (in green). Contacts are defined as amino acids within 8 Å in the 1ecm PDB structure. Experimentally sensitive positions obtained as in Fig. 6. When not visible, the blue curve is under the green curve. While both high and low regularization results are good at inferring contacting positions, it can be concluded that the contacts predicted in the high regularization result are distinct from the ones predicted by the low regularization result since they have a strong overlap with the experimentally sensitive position pairs, whereas low regularization contacts do not.

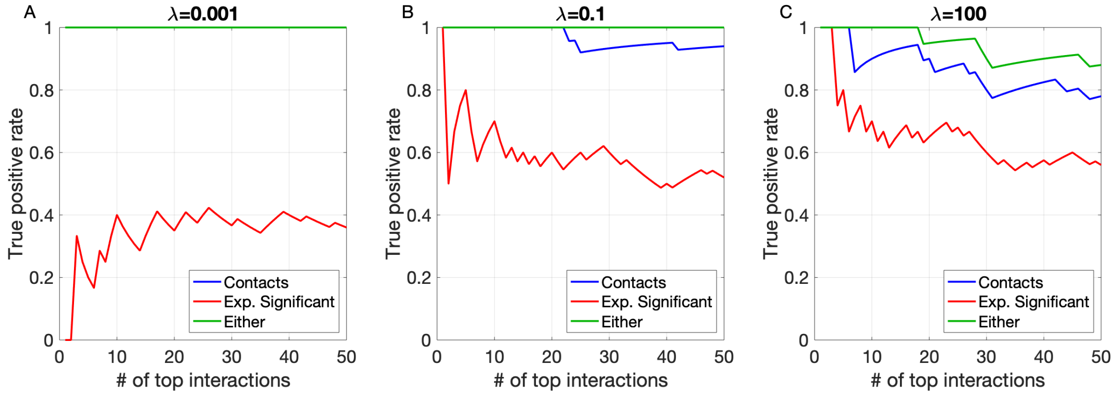


**Fig. S8.** Statistics over top coupled pairs, ranked by $\|\hat{J}_{ij}\|_{\mathrm{APC}}$, based on the APC defined in Eq. (S5) – **(A,B,C)** For three values of the regularization parameter $\lambda_J$, fraction of top $\|\hat{J}_{ij}\|_{\mathrm{APC}}$ pairs that are contact position pairs (in blue), experimentally sensitive pairs (in red) or in either one of these groups (in green). When compared to Fig. S7 we see that while applying APC greatly increases the predictive power for contacts, it is not obvious that this is the case for cooperative units.
